# Search for early pancreatic cancer blood biomarkers in five European prospective population biobanks using metabolomics

**DOI:** 10.1101/543686

**Authors:** Jesse Fest, Lisanne S. Vijfhuizen, Jelle J. Goeman, Olga Veth, Anni Joensuu, Markus Perola, Satu Männistö, Eivind Ness-Jensen, Kristian Hveem, Toomas Haller, Neeme Tonisson, Kairit Mikkel, Andres Metspalu, Cornelia M. van Duijn, Arfan Ikram, Bruno H. Stricker, Rikje Ruiter, Casper H.J. van Eijck, Gertjan B. van Ommen, Peter A.C. ’t Hoen

## Abstract

**Background and aim:** Most patients with pancreatic cancer present with advanced disease and die within the first year after diagnosis. Predictive biomarkers that signal the presence of pancreatic cancer in an early stage are desperately needed. We aimed to identify new and validate previously found plasma metabolomic biomarkers associated with early stages of pancreatic cancer.

**Methods:** The low incidence rate complicates prospective biomarker studies. Here, we took advantage of the availability of biobanked samples from five large population cohorts (HUNT2, HUNT3, FINRISK, Estonian biobank, Rotterdam Study) and identified prediagnostic blood samples from individuals who were to receive a diagnosis of pancreatic cancer between one month and seventeen years after blood sampling, and compared these with age- and gender-matched controls from the same cohorts. We applied ^1^H-NMR-based metabolomics on the Nightingale platform on these samples and applied logistic regression to assess the predictive value of individual metabolite concentrations, with gender, age, body mass index, smoking status, type 2 diabetes mellitus status, fasting status, and cohort as covariates.

**Results:** After quality assessment, we retained 356 cases and 887 controls. We identified two interesting hits, glutamine (p=0.011) and histidine (p=0.012), and obtained Westfall-Young family-wise error rate adjusted p-values of 0.43 for both. Stratification in quintiles showed a 1.5x elevated risk for the lowest 20% of glutamine and a 2.2x increased risk for the lowest 20% of histidine. Stratification by time to diagnosis (<2 years, 2-5 years, >5 years) suggested glutamine to be involved in an earlier process, tapering out closer to onset, and histidine in a process closer to the actual onset. Lasso-penalized logistic regression showed a slight improvement of the area under the Receiver Operator Curves when including glutamine and histidine in the model. Finally, our data did not support the earlier identified branched-chain amino acids as potential biomarkers for pancreatic cancer in several American cohorts.

**Conclusion:** While identifying glutamine and histidine as early biomarkers of potential biological interest, our results imply that a study at this scale does not yield metabolomic biomarkers with sufficient predictive value to be clinically useful *per se* as prognostic biomarkers.

## Introduction

Pancreatic cancer is one of the most lethal cancers worldwide and increasingly common(1-3). Most patients present with advanced and thus incurable disease and die within a year of the initial diagnosis(3,4). There is an imminent need to identify these patients earlier in the disease process, as patients with resectable, non-metastatic cancer can potentially be cured. For many cancers it takes several years for a local malignant lesion to progress to fully metastasized disease, and pancreatic cancer is no exception(5). Thus, there should be a window of opportunity for timely detection and intervention. Unfortunately, for early, presymptomatic pancreatic cancer currently no specific biomarkers are available. The identification of predictive biomarkers is complicated by the low incidence rate of the disease, estimated at 7-12 cases per 100,000 adult person years in the Western European population(6,7).

It is well known that the development and progression of pancreatic cancer are associated with alterations in systemic metabolism. Patients may present with glucose intolerance, anorexia and severe weight loss(3,8). In line with this, circulating metabolites have been proposed as a potentially useful screening tool in pancreatic cancer(9-16). The study by Mayers et al.(11) stood out from other metabolomic biomarker studies, as they analyzed blood samples taken two to more than ten years prior to diagnosis. They found an elevation of circulating branched-chain amino acids as an early event in the development of pancreatic cancer(11).

Considering these preliminary predictive metabolomics biomarkers as promising, we set out to replicate these findings independently in five large European population cohorts and find additional biomarkers, using a different platform, proton nuclear magnetic resonance (^1^H-NMR) instead of liquid chromatography followed by mass spectrometry (LC-MS).

## Methods

### Study Population

Our study population consisted of pancreatic cancer cases and controls, drawn from five national European cohorts, collaborating in the Biobanking and BioMolecular resources Research Infrastructure Large Population Cohorts (BBMRI-LPC; www.bbmri-lpc-biobanks.eu) and the cross-infrastructure project CORBEL (www.corbel-project.eu): the Estonian Genome Center of the University of Tartu study (EGCUT), the FINRISK Study (FR), the Nord-Trøndelag health study (HUNT2 and HUNT3), and the Rotterdam Study (RS).

EGCUT is a volunteer-based sample of the Estonian resident adult population aged 18 years and above, started in 1999 and currently has close to 52,000 participants(17).

FINRISK was initiated in 1972 and includes a collection of cross-sectional surveys in the adult (25 to 74-year-old) permanent residents of selected geographical areas of Finland. Altogether, FINRISK had nine cross sectional surveys performed every fifth year by the National Institute for Health and Welfare, including a total of 101,451 invitees(18). Participants in this study were selected from the FINRISK1997, 2002 and 2007 surveys. There are no re-examinations except for occasional persons who were selected to more than one independent survey by chance. Follow-up is carried out through record linkages to national administrative registers (such as the Causes of Death Register and Cancer Register), by using a unique personal identity code(19).

HUNT includes repeated surveys of a large population-based cohort in Norway. Data from 116,044 individuals aged 20 years and older from HUNT2 (1995-1997, n=65,237) and HUNT3 (2006-2008, n=50,807) were used in this study. Individuals who participated in both HUNT2 and HUNT3 were only included as part of HUNT3. Similar to FINRISK, follow-up is carried out through record linkages to national administrative registers (such as the Causes of Death Register and Cancer Register), by using a unique personal identity code(20).

The RS is an ongoing, population-based cohort study in a suburban area of Rotterdam, the Netherlands. It was initiated in 1989 and has enrolled 14,926 individuals of 45 years and older since then. Follow-up is carried out continuously(21,22).

All participants of the respective cohorts provided a written informed consent. The present study was approved by the local ethics committee of each study.

### Selection of cases and controls

We included incident pancreatic cancer cases, confirmed by pathology, and diagnosed after blood collection. Cases were identified through national cancer registries and through independent review of medical records. For diagnosis of pancreatic cancer, we used the ICD-10 C25.0 code. Deaths were ascertained through the national registries. We excluded cases that lived more than 5 years after diagnosis, to avoid false positive diagnoses(23-25).

For each case, we selected two (in RS one, in EGCUT four) random controls, matching on cohort, gender, age (± 2 years), and time of blood collection. Controls were those who were alive, without a diagnosis of pancreatic cancer at time of the case’s diagnosis date.

### Ascertainment of other covariates

The following covariate data were obtained from questionnaires and physical examination before blood collection: body mass index (BMI; kg/m^2^), smoking status (current/former/never), type 2 diabetes mellitus (T2DM) status, and fasting status (<4h/4-8h/>8h).

### Metabolite profiling and quality control

Metabolites were quantified from EDTA-plasma (EGCUT) or serum (HUNT2, HUNT3, FR, RS) samples using a high-throughput ^1^H-NMR metabolomics platform (Nightingale Health, Helsinki, Finland; https://nightingalehealth.com/). This platform provides simultaneous quantification of 147 individual metabolites and 79 metabolite ratios, e.g. routine lipids, lipoprotein subclass profiling with lipid concentrations within 14 subclasses, esterified fatty acid composition, and various low-molecular metabolites including amino acids, ketone bodies, and gluconeogenesis-related metabolites in molar concentration units. Details of the experimentation and applications of the platform have been described previously(26).

Metabolite measures that failed quality control (in particular for glutamine, pyruvate, glycerol, hydroxybutyrate, and acetate) were excluded from the analysis on a per-individual basis. One metabolite measure (glycerol) with >10% missing values was excluded entirely, resulting in a final number of 146 metabolite measures and 79 ratios. Outliers (>5 SD) were removed in concordance with previous research in this field(27).

### Statistical analysis

Differences in baseline characteristics between cases and controls were assessed for each cohort separately using Student’s t-tests and ANOVAs.

Metabolite measurements were raised by one to allow log-transformation. Thereafter all metabolite values were log-transformed and scaled to obtain unit standard deviation for each cohort. They were included as continuous variables in logistics regression models and adjusted for matching factors (gender and age, minimally adjusted model). In our main model on the pooled data from all the cohorts, we additionally adjusted for BMI, smoking status, T2DM status, fasting status, and cohort. P-values were corrected for multiple testing using Westfall and Young’s family-wise error rate, an appropriate method given the strong correlations between the measurements of the different metabolites(28). To provide estimates of effect magnitude, significant metabolites were again examined in logistic regression models after categorization in quintiles. Quintiles were generated based on the metabolite values in controls only. Results were presented as odds ratios (ORs) and 95% confidence intervals (CIs).

As an alternative for the pooling of the data from the different cohorts, we also performed a logistic regression per cohort (with gender, age, BMI, smoking status, T2DM, and fasting status as covariates) and a subsequent meta-analysis. The obtained estimates for the metabolite measures and their standard errors were used in a random effects meta-analysis using the R-package meta 4.9.2(29). A random effects model was chosen to account for possible heterogeneity due to differences in disease assessment, sample processing, and sample collection between cohorts. Heterogeneity was assessed using the I^2^-statistic and by visual inspection of forest plots. P-values from the meta-analysis were corrected for multiple-testing using Bonferroni-Holm.

### Lasso regression to evaluate additive effect of metabolomics biomarkers on top of clinical predictors

To select biomarkers with predictive value, we applied a 5-fold cross-validated penalized lasso regression with the penalized package version 0.9-51(30). The clinical covariates (gender, age, BMI, smoking status, T2DM status, fasting status, and cohort) were not penalized and thus always present in the model. We performed a stratified analysis, including all controls but only cases who developed pancreatic cancer within 2 years or within 5 years after blood sampling or including all cases. For the variable selection, the data were split randomly into a dataset for variable selection (70% of the data; with 35% for training and 35% for cross-validation) and a dataset for performance testing (30% of the data). We compared the performance of the null model (with only the clinical covariates) with the model that included the selected metabolites using an ordinary least squares regression model. The performance of the different model was assessed by evaluating the area under the receiver operator curve (AUC).

### General

Analyses were performed using the software packages meta 4.9-2, Penalized 0.9-51, Globaltest 5.24.0, InformationValue 1.2.3, ROCR 1.0-7, RColorBrewer 1.1-2, and ggplot2 3.0.0 for R Version 3.2.3.

## Results

### Study population and measurements

Cross-checking of the individuals in the five population cohorts included in this study with the national cancer registries enabled us to identify 444 prediagnostic samples from subjects who received diagnosis of pancreatic cancer between 1 month and 17 years after blood sampling (median 4.68 years). We subsequently selected 1,012 gender-and age-matched controls from the same cohorts (**Figure 1**). Baseline characteristics for all cohorts are shown in **Table 1**. Although baseline characteristics differed significantly between cohorts (in particular for gender, BMI, T2DM, and fasting status), cases and controls did not differ in these characteristics within a cohort. We reliably quantified 146 blood metabolites and 79 metabolite ratios. **Figure 1** shows the number of participants remaining after quality control and after assessment of the completeness of phenotype information in the different analyses performed.

**Figure 1.**
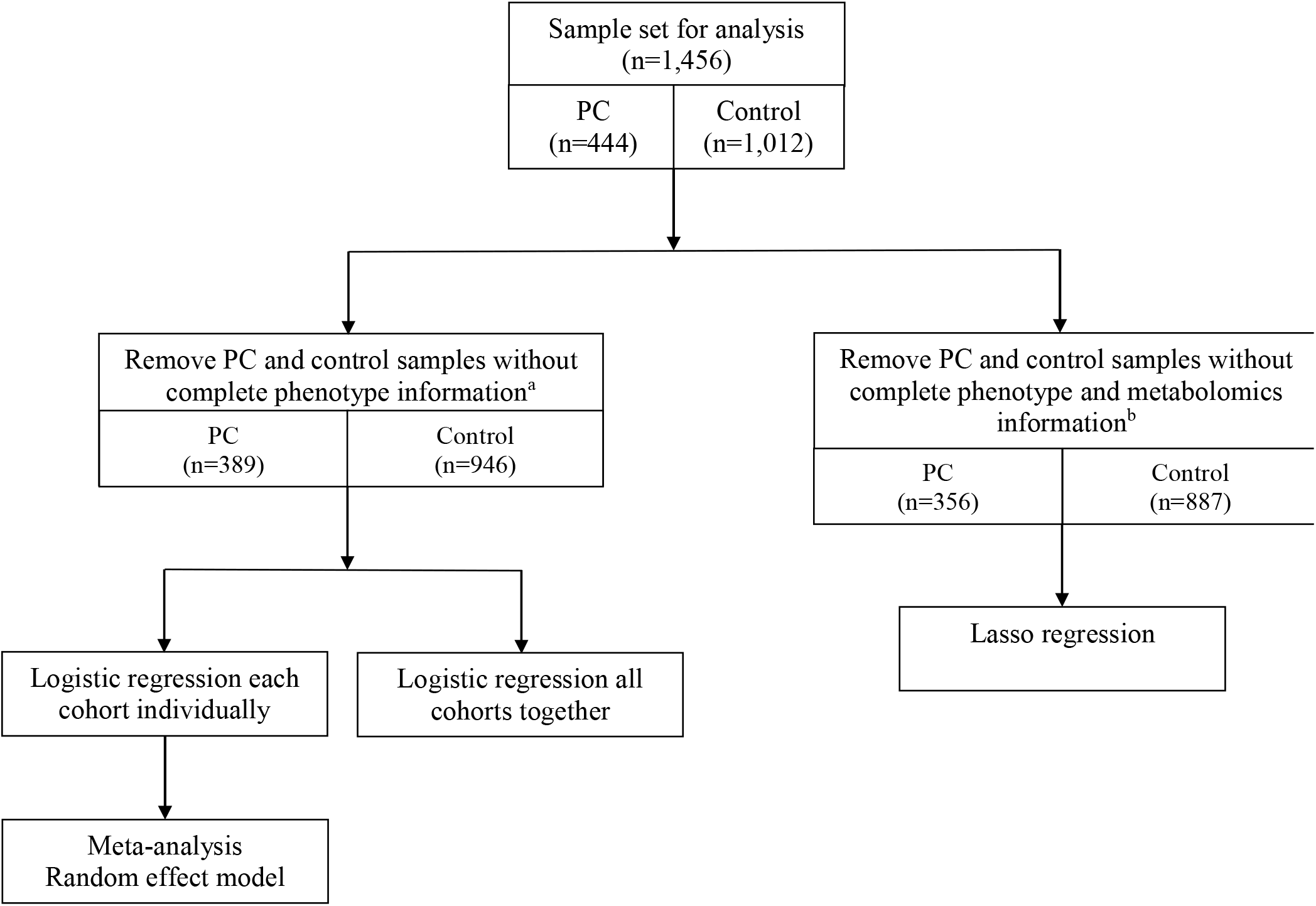
Schematic overview of the sample set used for data-analysis and the different data-analysis approaches performed in the current study. Footnotes: ^a^Any individual containing missing values in metabolomics measurements or phenotypically information were assumed to be missing at random, and removed from the dataset. ^b^Any individual containing missing values in phenotypically information were removed from the dataset. Abbreviation: PC - pancreatic cancer.

**Table 1:**
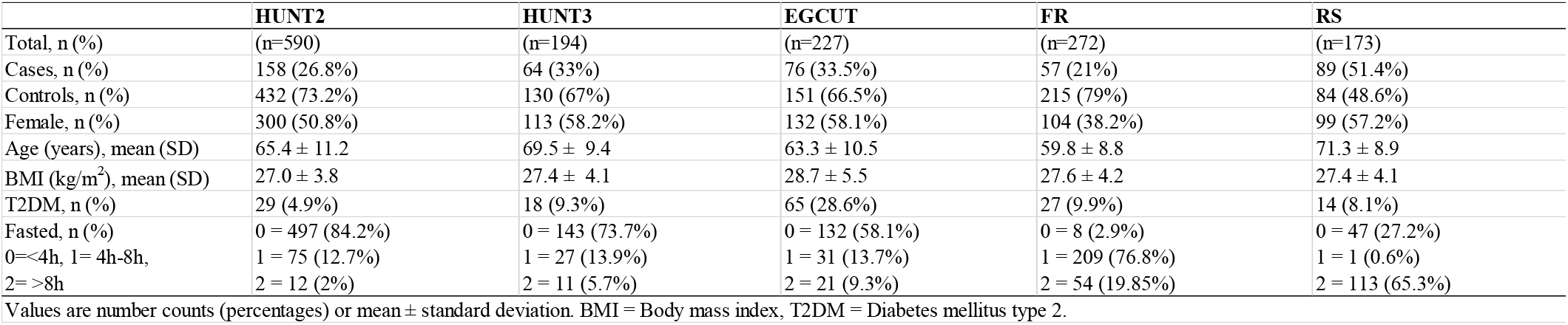
Baseline characteristics of samples.

|  | HUNT2 | HUNT3 | EGCUT | FR | RS |
| --- | --- | --- | --- | --- | --- |
| Total, n (%) | (n=590) | (n=194) | (n=227) | (n=272) | (n=173) |
| Cases, n (%) | 158 (26.8%) | 64 (33%) | 76 (33.5%) | 57 (21%) | 89 (51.4%) |
| Controls, n (%) | 432 (73.2%) | 130 (67%) | 151 (66.5%) | 215 (79%) | 84 (48.6%) |
| Female, n (%) | 300 (50.8%) | 113 (58.2%) | 132 (58.1%) | 104 (38.2%) | 99 (57.2%) |
| Age (years), mean (SD) | 65.4 ± 11.2 | 69.5 ± 9.4 | 63.3 ± 10.5 | 59.8 ± 8.8 | 71.3 ± 8.9 |
| BMI (kg/m<sup>2</sup>), mean (SD) | 27.0 ± 3.8 | 27.4 ± 4.1 | 28.7 ± 5.5 | 27.6 ± 4.2 | 27.4 ± 4.1 |
| T2DM, n (%) | 29 (4.9%) | 18 (9.3%) | 65 (28.6%) | 27 (9.9%) | 14 (8.1%) |
| Fasted, n (%) | 0 = 497 (84.2%) | 0 = 143 (73.7%) | 0 = 132 (58.1%) | 0 = 8 (2.9%) | 0 = 47 (27.2%) |
| 0 < 4h, 1 = 4h-8h, | 1 = 75 (12.7%) | 1 = 27 (13.9%) | 1 = 31 (13.7%) | 1 = 209 (76.8%) | 1 = 1 (0.6%) |
| 2 = > 8h | 2 = 12 (2%) | 2 = 11 (5.7%) | 2 = 21 (9.3%) | 2 = 54 (19.85%) | 2 = 113 (65.3%) |

Values are number counts (percentages) or mean ± standard deviation. BMI = Body mass index, T2DM = Diabetes mellitus type 2.

### Single-metabolite logistic regression

To identify metabolite biomarkers potentially associated with future pancreatic cancer diagnosis, we performed a separate logistic regression for each metabolite measured. In our primary model, we adjusted for the following covariates: gender, age, BMI, smoking status, T2DM status, fasting status, and cohort. The results of our top metabolites are presented in **Table 2**. Full data are provided in the **Supplementary Table 1**. Two metabolites demonstrated lower blood levels in cases than in controls and nominal significance: glutamine (p=0.012) and histidine (p=0.011). They were not significant after adjustment for multiple testing (Westfall-Young family-wise error rate adjusted p-value 0.43 for both metabolites). A closer inspection of the levels of glutamine and histidine, revealed that the differences were consistently observed across cohorts (**Figure 2A,E**), except for glutamine in RS and histidine in FR. Glutamine levels were lower in both non-diabetics and diabetics, whereas lower histidine levels were mainly observed in pancreatic cancer that were also diagnosed with T2DM (**Figure 2B**). Histidine levels were lower in individuals who developed pancreatic cancer within 2 years after blood sampling, whereas glutamine levels were decreased longer before diagnosis (**Figure 2C,G**). Histidine levels were lower in both fasting and non-fasting individuals (**Figure 2H**), whereas the effect of fasting status on glutamine levels is difficult to ascertain given the differences between cohorts in fasting status (**Figure 2D)**. The branched chain amino acids, leucine, valine, and isoleucine, reported earlier by Mayers et al.(11), were not different between cases and controls (unadjusted p-values of 0.75, 0.94, and 0.61, respectively).

**Table 2:**
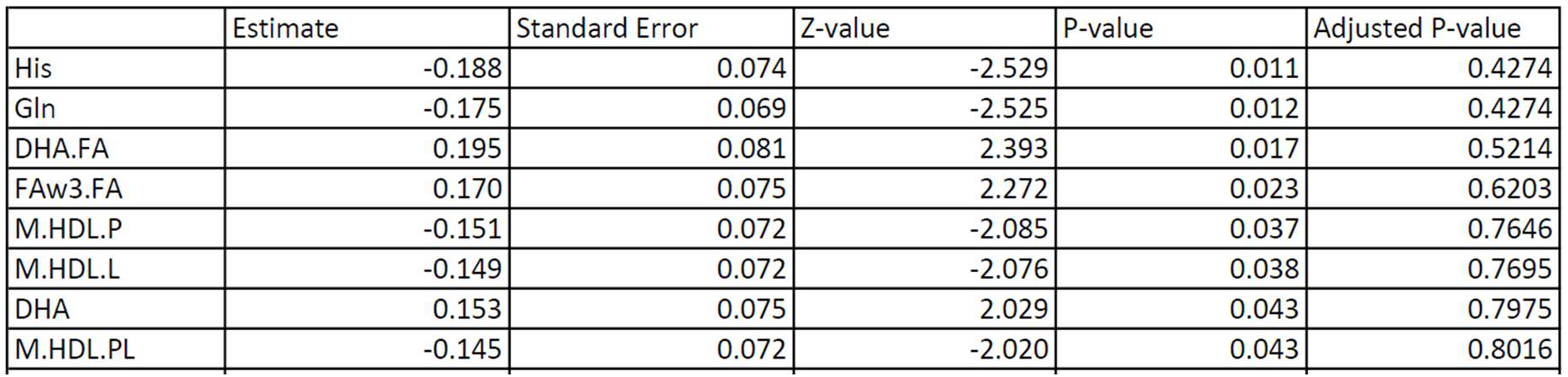
Top hits from logistic regression analysis. Logistic regression with single metabolite measure, BMI, smoking status, T2DM status, fasting status, and cohort as covariates. Explanation of the abbreviations of the metabolites. His: histidine; Gln: glutamine; DHA.FA: ratio of docosahexaenoic acid to all fatty acids; FAw3.FA: ratio of omega-3 fatty acids to total acids; M.HDL.P: concentration of medium HDL particles; M.HDL.L: total lipids in medium-sized HDL particles; DHA: docosahexaenoic acid; M.HDL.PL: phospholipids in medium-sized HDL particles

**Figure 2.**
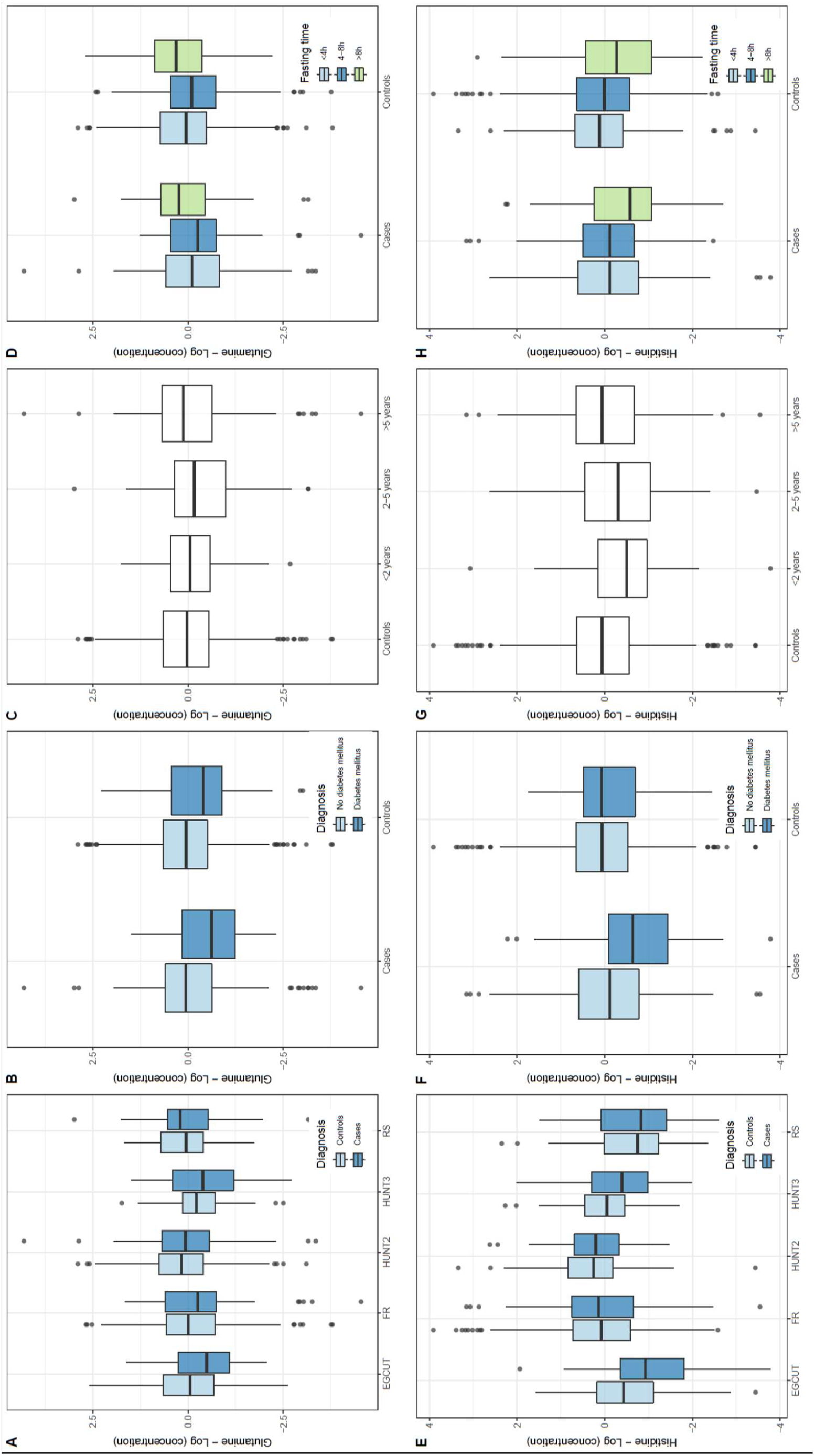
Concentrations (logarithmic scale) of glutamine (panel **A-D**) and histidine (panel **E-H**) in the blood circulation in controls and cases, *i.e.* those individuals who developed pancreatic cancer within a time window after blood sampling. **A,E**: Distribution of the concentrations of controls (light blue) and cases (dark blue) in the different cohorts analyzed (EGCUT, FR, HUNT2, HUNT3, RS). **B,F**: Distribution of concentrations in non-diabetics (light blue) and individuals diagnosed with T2DM (dark blue). **C,G**: Distribution of concentrations in controls and cases sampled within 2 years before diagnosis, between 2-5 years before diagnosis, and more than 5 years before diagnosis. **D,H:** Distribution of concentrations in non-fasting individuals (light blue), individuals who had a meal between 4 and 8 hours before blood draw (dark blue) and fasting individuals (green, last meal was more than 8 hours before blood draw). Box plots reflect the distribution of the concentrations in individual samples: the middle quartiles (25-75% of the data points are in the boxes), the horizontal band the median value, the lower whiskers represent the data points up to 1.5*the interquartile range (IQR) below the 25%, the upper whiskers represent the data points up to 1.5*IQR above the 75%; the data points outside these ranges are plotted as individual data points.

The results above were recapitulated in a minimally adjusted model, only corrected for gender and age (**Supplementary Table 2**). Glutamine and histidine were still among the top hits, with p-values of 0.0063 and 0.00045 (not adjusted for multiple testing), respectively.

To further address potential cohort differences, we performed a meta-analysis on the beta coefficients from the logistic regression models that were applied per cohort. The results are summarized in **Table 3** and provided in full in **Supplementary Table 3**. The results from the meta-analysis corroborated our findings on the pooled data, with lower glutamine levels seen for all cohorts (unadjusted p-value 0.0040), but most prominently in HUNT3 (**Figure 3A**), and lower histidine levels mostly in HUNT3 and EGCUT (unadjusted p-value 0.0022) (**Figure 3B**, with similar trends in other cohorts and evidence for significant heterogeneity between cohorts). The meta-analysis provided some evidence for the involvement of omega-3 fatty acids (FAw3, including docosahexaenoic acid (DHA)) and high density lipoproteins (HDL).

**Table 3:** Top hits from meta-analysis. Meta-analysis across the five cohorts of logistic regression results with single metabolite measure, BMI, smoking status, T2DM status, and fasting status as covariates. B is effect size; CI is confidence interval; P-value is Holm-Bonferroni adjusted p-value. I^2^ is the statistic used for heterogeneity between cohorts. Explanation of the abbreviations of the metabolites. Gln: glutamine; DHA.FA: ratio of docosahexaenoic acid to total fatty acids; M.HDL.PL: phospholipids in medium-sized HDL particles; M.HDL.P: concentration of medium-sized HDL particles; M.HDL.L: total lipids in medium-sized HDL particles; FAw3.FA: ratio of omega-3 fatty acids to total acids; His: histidine; M.HDL.FC: free cholesterol in medium-sized HDL particles; M.HDL.C: total cholesterol in medium-sized HDL particles; M.HDL.CE: cholesterol esters in medium-sized HDL particles; DHA: docosahexaenoic acid.

| Metabolite abbreviation | B | CI | Unadjusted P-value | P-value | I<sup>2</sup> (%) |
| --- | --- | --- | --- | --- | --- |
| Gln | -0.19538 | -0.33:-0.06 | 0.004 | 0.9037 | 0 |
| DHA.FA | 0.17259 | 0.04:0.3 | 0.0083 | 1 | 0 |
| M.HDL.PL | -0.17905 | -0.32:-0.04 | 0.0103 | 1 | 0 |
| M.HDL.P | -0.17856 | -0.32:-0.04 | 0.0104 | 1 | 0 |
| M.HDL.L | -0.17732 | -0.31:-0.04 | 0.0104 | 1 | 0 |
| FAw3.FA | 0.16222 | 0.03:0.29 | 0.0126 | 1 | 0 |
| His | -0.25164 | -0.46:-0.05 | 0.0156 | 1 | 0.53 |
| M.HDL.FC | -0.15636 | -0.29:-0.02 | 0.0251 | 1 | 0 |
| M.HDL.C | -0.15174 | -0.29:-0.02 | 0.0267 | 1 | 0 |
| M.HDL.CE | -0.14723 | -0.28:-0.01 | 0.0306 | 1 | 0 |
| DHA | 0.13222 | 0:0.26 | 0.0438 | 1 | 0 |

**Figure 3.**
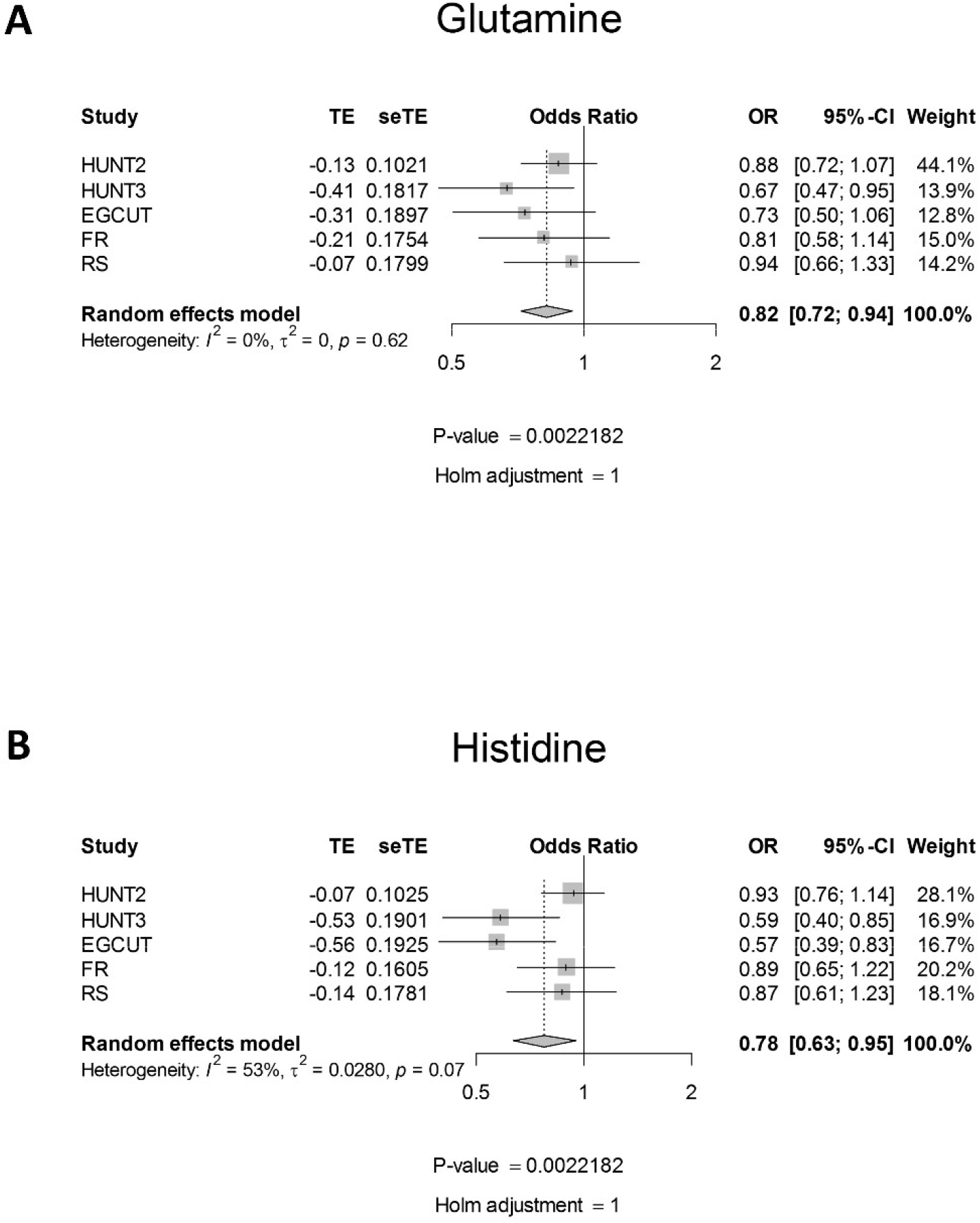
Forest plots from random-effects meta-analysis across different cohorts for glutamine (panel **A**) and histidine (panel **B**). The meta-analysis was performed on the beta coefficients and standard deviations from the logistic regressions run for each cohort separately. In the logistic regression, pancreatic cancer status was modeled as a function of log-transformed and standardized metabolite concentration, gender, age, BMI, smoking status, T2DM and fasting status. Shown are the estimated effect size, the standard error on this estimate, the estimated odds ratio and the confidence interval on this ratio, the weight of the individual cohort on the calculation of the final estimate, the heterogeneity measure (modeling differences between cohorts), and the unadjusted and Bonferroni-Holm corrected p-values for the respective metabolite.

To provide a better understanding of the risks associated with lower glutamine or histidine levels, we stratified the cohorts in quintiles based on the glutamine or histidine levels in controls. Individuals within the lowest 20% of glutamine levels ran a 1.5 times elevated risk of pancreatic cancer and individuals within the lowest 20% of histidine levels ran a 2.2 times elevated risk of pancreatic cancer (**Table 4**).

**Table 4:** Odds ratios for developing pancreatic cancer in different glutamine (top) and histidine (bottom) strata.

| Gln | Based on control data | N (controls) | N (cases) | OR | CI 5% | CI 95% | Pvalue |
| --- | --- | --- | --- | --- | --- | --- | --- |
| 0% | 0.269 | 180 | 94 | 1 | - | - |  |
| 20% | 0.4538 | 176 | 66 | 0.72 | 0.49 | 1.05 | 0.0852 |
| 40% | 0.487 | 177 | 62 | 0.67 | 0.46 | 0.98 | 0.0404 |
| 60% | 0.5157 | 176 | 71 | 0.77 | 0.53 | 1.12 | 0.1734 |
| 80% | 0.55358 | 178 | 62 | 0.66 | 0.46 | 0.98 | 0.0376 |

| His |  | N (controls) | N (cases) | OR | CI 5% | CI 95% | Pvalue |
| --- | --- | --- | --- | --- | --- | --- | --- |
| 0% | 0.03927 | 178 | 110 | 1 | - | - |  |
| 20% | 0.060498 | 177 | 71 | 0.65 | 0.45 | 0.93 | 0.0199 |
| 40% | 0.064778 | 177 | 58 | 0.53 | 0.36 | 0.78 | 0.0011 |
| 60% | 0.068174 | 177 | 66 | 0.6 | 0.42 | 0.87 | 0.0073 |
| 80% | 0.072638 | 178 | 51 | 0.46 | 0.31 | 0.69 | 0.0001 |

### Lasso regression

Lasso regression was used to evaluate the additional predictive effect of metabolomics biomarkers over clinical predictors. The performance of a reference (null) model, in which only the clinical covariates were used for prediction, was compared with an alternative model, in which metabolites selected by the lasso regression were added to the model. The cases were stratified according to the time until diagnosis (up to 2 years, up to 5 years and all cases without temporal constraint). In the model with cases up to 2 years until diagnosis, the lasso regression selected medium very low-density lipoprotein (VLDL), total unsaturated fatty acids and saturated fatty acids to be included in the model (**Table 5**), but it did not affect the performance on the 30% of the data that were unseen during the selection of the metabolites. In the model with cases up to 5 years until diagnosis, the lasso regression model selected small VLDL and glutamine (consistent with the prominent decrease of glutamine levels in cases between 2 and 5 years before diagnosis) (**Table 5**). The performance of the alternative model increased slightly for both the training (AUC = 0.72 vs 0.71 for the null model, **Figure 4A**) and the validation set (AUC=0.64 vs 0.62 for the null model, **Figure 4B**). In the model with all cases included, more metabolites were selected (**Table 5**), but the performance of the model including the metabolites on both training and validation set (AUC of 0.68 and 0.62, respectively) was worse than for the model with cases up to 5 years.

**Table 5:**
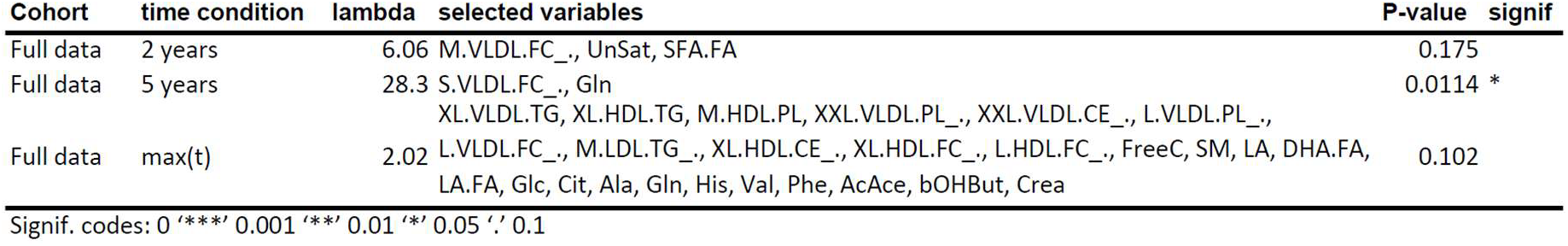
Variables selected by the lasso regression. 1) The results of the cross-validated Lasso-penalized logistic regression for full dataset. For each regression the penalty parameter (λ) and the selected covariates (separated by commas) are given. For every model where metabolites were selected, the significance of the presence of all the selected metabolites in the model compared to the model without presence of metabolites is tested in a global test, and its p-value is given here. Note that the p-value is only for the metabolites, not for the clinical covariates. 2) Abbreviations of metabolites mentioned: M.VLDL.FC_.: free cholesterol to total lipids ratio in medium VLDL; UnSat: estimated degree of unsaturation; SFA.FA: ratio of saturated fatty acids to total fatty acids; S.VLDL.FC_.: free cholesterol to total lipids ratio in small VLDL; Gln: glutamine; XL.VLDL.TG: triglycerides in extra large VLDL particles; XL.HDL.TG: triglycerides in very large HDL; M.HDL.PL: phospholipids in medium-sized HDL; XXL.VLDL.PL_.: phospholipids to total lipids ratio in chylomicrons and extremely large VLDL; XXL.VLDL.CE_.: cholesterol esters to total lipids ratio in chylomicrons and extremely large VLDL; L.VLDL.PL_.phospholipids to total lipids ratio in large VLDL; L.VLDL.FC_.: free cholesterol to total lipids ratio in large VLDL; M.LDL.TG_.: triglycerides to total lipids ratio in medium LDL; XL.HDL.CE_.: cholesterol ester to total lipids ratio in very large HDL; XL.HDL.FC_.: free cholesterol to total lipids ratio in very large HDL; L.HDL.FC_.: free cholesterol to total lipids ratio in large HDL; FreeC: free cholesterol; SM: sphingomyelins; LA: linoleic acid; DHA.FA: ratio of docosahexaenoic acid to total fatty acids; LA.FA: ratio of linoleic acid to total fatty acids; Glc: glucose; Cit: citrate; Ala: alanine; Gln: glutamine; His: histidine; Val: valine; Phe: phenylalanine; AcAce: acetoacetate; bOHBut: 3-hydroxybutyrate; Crea: creatinine

**Figure 4.**
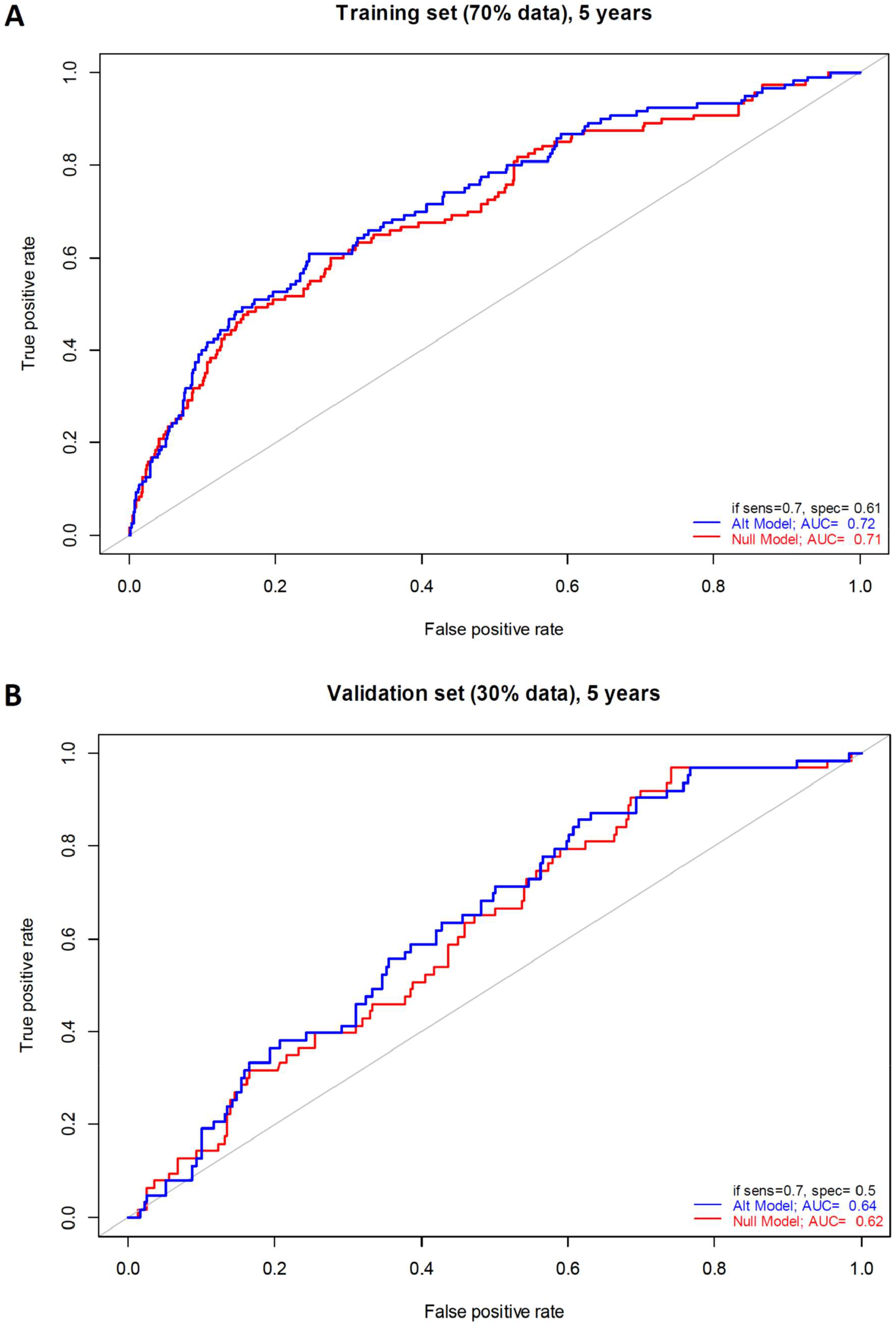
Receiver operator curves for classification of pancreatic cancer cases (sampled up to 5 years before diagnosis) and controls for training (70% of all individuals, panel **A**) and performance testing (30% of all individuals unseen during the variable selection, panel **B**) set. In red, the null model in which only the clinical covariates (gender, age, BMI, smoking status, T2DM, and fasting status) were included in the regression. In blue, the alternative model where the metabolites selected by the lasso regression were included in addition to the clinical covariates. The receiver operating curve AUCs are indicated, as well as the specificity (1 - false positive rate) at 70% sensitivity.

## Discussion

Pancreatic cancer is usually diagnosed in an advanced stage of the disease, resulting in a poor prognosis. Most pancreatic cancer biomarker studies executed until today(9,10,13-16) collected samples at the time of diagnosis or even later, and therefore have limited clinical utility. However, they may provide insight in the pathophysiology of the disease. The setup of our study allowed for the identification of prospective biomarkers, and made efficient use of the large scale biobanking infrastructure in Europe (BBMRI-LPC program).

We identified two of these potentially, prognostic biomarkers, glutamine and histidine, while noting that the clinical utility of these biomarkers is currently low. They did not reach significance after multiple testing correction was taken into account and it is therefore possible that these hits are false positives. The increased risk of pancreatic cancer associated with low levels of glutamine and histidine was only 1.5 – 2.2-fold and do not add much in terms of predictive potential to well-known risk factors for pancreatic cancer such as age, smoking and T2DM. However, also earlier studies provided evidence for alterations in glutamine and histidine in pancreatic cancer (10,15,16) suggesting that these may indeed be associated with pancreatic cancer-associated changes in metabolism. In the largest study by Fukutake *et al.*(15) (N=360 vs 8372), histidine was found particularly low in patients with resectable disease stage 0-IIB. This group of patients in a relatively early state of the disease is likely most similar to our group of individuals whom were diagnosed in less than two years after blood sampling and had the lowest histidine levels of all cases. Also in other cancer-related studies, negative correlations between histidine levels and cancer incidence and/or cancer-associated mortality were observed(31-33). Remarkably, a recent report demonstrated also lower efficacy of cancer treatment in individuals with low histidine levels, and suggested histidine supplementation to enhance the efficacy of methotrexate treatment in leukemia(34).

One of the reasons why changes in metabolites like glutamine and histidine are difficult to detect is that the concentrations of these metabolites are relatively high, and that local events like a pancreatic tumor, contribute only little to the overall pool of these metabolites. Other metabolites may be more specific to the metabolism in the pancreas and may show more prominent changes. This type of metabolites require broader metabolomic screening than the Nightingale platform provides. While having superior robustness and throughput and low cost, the range of metabolites measured on the Nightingale platform is limited to amino acids, other polar metabolites, and a large range of lipid and lipoprotein classes. Our study calls for the use of complementary biomarker platforms on these samples, and suggests to limit the sampling to within 5 years before diagnosis and not beyond.

Our study was not able to replicate the findings from the single study with a design and sample size comparable to ours(11). This study identified the branched chain amino acids valine, leucine, and isoleucine as potential prognostic biomarkers for pancreatic cancer. We did not find any difference between cases and controls for these amino acids nor were our top metabolites identified in this earlier study. This may be a reflection of the limited power of both studies for the discovery of small changes observed for these metabolites. However, we did not even observe trends in the same directions. Differences in the measurement platforms (^1^H-NMR vs. LC-MS) may play a role, but the different amino acids can robustly be measured by both. It is equally plausible that the differences are due to differences in the studied populations or confounding factors, which were not or were incompletely corrected for in the statistical model, such as nutrition.

In conclusion, our study lends initial support to the existence of metabolic alterations in early pancreatic cancer development, highlighting glutamine and histidine as metabolites of interest, but also underscores the challenges to find robust, prognostic biomarkers for rare disorders. To address this, larger studies are needed, including more metabolites with lower concentrations, and/or integrated studies at multiple -omics levels.

## Acknowledgements

Tony Wilkes and Abdelhak Chahid (Leiden University) are gratefully acknowledged for their contribution to the analysis presented in the manuscript. Annika Wolin is thanked for her contribution to the FINRISK sample selection. We would like to thank Peter Würtz (Nightingale) for very useful comments on the study. This work was carried out on the Dutch national e-infrastructure with the support of SURF Cooperative.

The Nord-Trøndelag Health Study (The HUNT Study) is a collaboration between HUNT Research Centre (Faculty of Medicine and Health Sciences, Norwegian University of Science and Technology NTNU), Trøndelag County Council, Central Norway Regional Health Authority, and the Norwegian Institute of Public Health.

The study has used data from the Cancer Registry of Norway. The interpretation and reporting of these data are the sole responsibility of the authors, and no endorsement by the Cancer Registry of Norway is intended nor should be inferred.

## Author contribution

MP, ENJ, KH, AM, CMvD, GJBvO and PACtH jointly designed the study. JF, JJG, PACtH, TH, NT, BHS, RR, CHJvE designed the analysis plan and statistical framework. JF, LSV, PACtH, AJ performed the analyses. MP, SM, ENJ, KH, KM, AM, CMvD, AI contributed samples. JF, LS, PACtH drafted the manuscript. All co-authors reviewed and edited the manuscript.

